# From divergence to contact: demographic history and genomic context shape introgression across independent damselfly hybrid zones

**DOI:** 10.64898/2026.04.09.717498

**Authors:** Miguel Stand-Pérez, Luis Rodrigo Arce-Valdés, Jesús Ernesto Ordaz-Morales, Janne Swaegers, Jesús Ramsés Chávez-Ríos, Carla Gutiérrez-Rodríguez, Enrique Ibarra-Laclette, Bengt Hansson, Fernanda Baena-Díaz, Rosa Ana Sánchez-Guillén

## Abstract

Hybridisation outcomes often vary across space and time, yet the relative roles of demographic history and genomic architecture in shaping introgression remain unclear. Here, we investigate three replicated hybrid zones between the damselflies *Ischnura elegans* and *Ischnura graellsii* across Spain to test whether genomic introgression patterns are repeatable across independently formed zones. Using genome-wide data, we combined demographic modelling, genomic cline, and functional annotation of introgressed loci. Demographic inference supported three independent secondary contact events of different ages: The South-east zone forming first (∼207 years ago), followed by the North-west (∼73.5 years) and North-central (∼33 years) hybrid zones. Despite similar cline steepness across autosomes, asymmetric gene flow from *I. graellsii* into *I. elegans* was observed, with a low overlap of introgressed loci between zones. These loci were mainly associated with broad regulatory and transport-related functions in both hybrid zones, indicating repeatability at the level of gene function rather than gene identity. In contrast, the X chromosome showed steeper clines, suggesting strong intrinsic genomic constraints. Together, demographic history explains geographic heterogeneity in introgression, whereas chromosome architecture imposes consistent constraints. These findings highlight how replicated hybrid zones can disentangle contingent versus repeatable genomic responses during early stages of speciation with gene flow.

## INTRODUCTION

Hybridisation is widely recognised as a dynamic evolutionary process capable of generating diverse outcomes ranging from genetic swamping, species replacement, reinforcement, hybrid speciation, stable hybrid zones, or adaptive introgression (Abbott et al., 2013). However, these outcomes are rarely uniform, even within the same pair of species (e.g. Arce-Valdés, Swaegers, et al., 2025). Growing evidence show that the consequences of hybridisation depend not only on intrinsic factors, but also on the demographic and environmental context in which secondary contact occurs (Peñalba et al., 2024). Understanding why similar species interactions lead to divergent genomic and evolutionary trajectories remains a central challenge in evolutionary biology.

One key source of variation arises from differences in demographic history and the timing of secondary contact. Hybrid zones formed at different times may experience contrasting reproductive barriers, effective population sizes, genetic incompatibilities, and ecological conditions, all of which influence the strength and direction of introgression (Harrison & Larson, 2014, 2016). Distributional range shifts driven by climate change or post-glacial warming have repeatedly generated scenarios in which closely related taxa come into contact under heterogeneous demographic contexts, creating natural laboratories for studying speciation with gene flow (Taylor et al., 2015). Yet most studies focus on single hybrid zones, limiting our ability to disentangle repeatable genomic responses from historically contingent outcomes.

In addition to demographic processes, genomic architecture plays an important role in shaping introgression patterns. Sex chromosomes and some genomic regions associated with reproductive isolation often show reduced permeability to gene flow (Dobzhansky, 1936; Fraïsse & Sachdeva, 2021), suggesting that intrinsic genetic barriers can impose predictable constraints across hybrid systems. At the same time, widespread variation in autosomal introgression among populations indicates that introgression patterns are not solely determined by species-specific differences but emerge from interactions between genome architecture and local demographic or ecological conditions (Abbott et al., 2016). This dual role of intrinsic genomic constraints and extrinsic historical context raises the issue of the extent to which introgression patterns are repeatable across independently formed hybrid zones within the same species pair.

Comparisons among multiple hybrid zones provide a powerful framework to investigate this issue. Spatially replicated zones differing in age, demographic context, and ecological conditions allow us to evaluate whether hybridisation produces consistent genomic signatures or whether outcomes diverge depending on historical contingencies (e.g. Arce-Valdés, Swaegers, et al., 2025; Harrison & Larson, 2016; Vines et al., 2003). Such systems align with the mosaic hybrid zone concept, in which geographically structured variation in selection, demography, and environmental context generates heterogeneous genomic landscapes of introgression (Rand & Harrison, 1989). Importantly, examining replicated contact zones within a single species pair offers a unique opportunity to separate lineage-specific genomic effects from broader demographic processes (Harrison & Larson, 2016). However, most previous evidence for repeatable patterns of introgression has been derived from interspecific comparisons, either across different species pairs or among independent hybrid systems (Edelman et al., 2019; Fontaine et al., 2015; Gompert et al., 2012; Martin et al., 2013; Taylor & Larson, 2019). This highlights the need for systems that allow such patterns to be evaluated within a single species pair across multiple hybrid zones.

A particularly useful framework for quantifying these patterns is the genomic cline theory (Barton, 1979), which can describe how introgression rates may vary across loci and between zones. Unlike traditional clines based on geography, genomic clines use hybrid indices or admixture proportions as independent variables (Szymura & Barton, 1991). This approach is especially powerful for detecting outlier loci whose introgression patterns deviates significantly from the genome-wide neutral expectation (Bailey, 2022; Fitzpatrick, 2013). Thus, the rate and pattern of introgression at a given locus can reveal whether it is linked to reduced hybrid fitness or, conversely, to adaptive or heterotic advantages (Gompert & Buerkle, 2011).

Our study system focuses on two closely related damselfly species that provide a rare opportunity to investigate recent divergence, range expansion, and hybridisation in real time. *Ischnura elegans* is a widespread Euro-Siberian species that has undergone rapid southward range expansion, whereas its sister species *I. graellsii* has a much narrower distribution, being largely restricted to the Iberian Peninsula and northern Africa (Ocharan-Larrondo, 1987). Over the last decades, and with increasing speed in recent years, *I. elegans* has expanded into the range of *I. graellsii* in Spain (Arce-Valdés, Swaegers, et al., 2025; Monetti et al., 2002; Wellenreuther et al., 2018) generating several geographically structured hybrid zones.

These hybrid zones differ markedly in their ecological and geographic context, the strength and asymmetry of reproductive barriers, patterns of introgression, and evolutionary outcomes (Arce-Valdés, Swaegers, et al., 2025). The South-east (SE) hybrid zone, located along the Mediterranean coast, represents the oldest documented zone and is characterised by introgression toward *I. elegans* and the progressive replacement of *I. graellsii* by *I. elegans* (Sánchez-Guillén et al., 2011; Wellenreuther et al., 2018). By contrast, the North-west (NW) hybrid zone is geographically isolated and exhibits extensive bidirectional introgression, reduced interspecific genetic differentiation, and reinforcement acting primarily on mechanical barriers (Arce-Valdés, Ballén-Guapacha, et al., 2025; Ballén-Guapacha et al., 2024; Sánchez-Guillén et al., 2012). Finally, the North-central (NC) hybrid zone is larger and currently expanding, showing predominantly unidirectional introgression toward *I. graellsii* and stronger postmating isolation involving both mechanical and fertility barriers (Arce-Valdés, Swaegers, et al., 2025; Sánchez-Guillén et al., 2011).

Across the three hybrid zones, genomic analyses reveal consistently reduced introgression on the X chromosome and an underrepresentation of males among admixed individuals, consistent with Haldane’s rule in an X0 sex-determination system (Swaegers, Sánchez-Guillén, Chauhan, et al., 2022). Together, these hybrid zones constitute a powerful natural laboratory for examining the repeatability and contingency of hybridisation outcomes, the interaction between range expansion and reinforcement, and the genomic architecture of reproductive isolation during the earliest stages of speciation. Beyond their role as a model for hybridization and reinforcement, recent studies have shown that the southward expansion of *I. elegans* has been accompanied by rapid ecological and physiological responses to warmer climatic conditions, including convergence in thermal tolerance and life-history traits toward the native sister species *I. graellsii* (Swaegers, Sánchez-Guillén, Carbonell, et al., 2022). These findings highlight the colonisation of warmer environments by *I. elegans* is reshaping the ecological context in which secondary contact and hybridisation occur.

Despite numerous studies conducted in this hybrid system, the origins of each hybrid zone remain unresolved as does the dual role that demographic conditions and genomic architecture may play in shaping the evolutionary consequences of hybridisation. For this reason, this study aims to investigate the demographic history and patterns of genomic introgression between *I. elegans* and *I. graellsii* across three hybrid zones in Spain. Specifically, we aim to (i) reconstruct the demographic history of this hybrid system by estimating the timing of the cessation of gene flow between species (origin of the allopatric distribution of the species), the formation of subsequent hybrid zones, and key demographic parameters. Given historical records of the southward expansion of *I. elegans*, we expect hybrid zones to have originated through independent and temporally staggered secondary contact events associated with this range expansion (Monetti et al., 2002; Wellenreuther et al., 2018; Arce-Valdés, Swaegers, et al., 2025). (ii) Model genomic clines within each hybrid zone to characterise patterns of genetic introgression. Based on previous evidence of asymmetric introgression and differences in reproductive barriers among zones, we predict that genomic clines will differ in the direction and magnitude of introgression among hybrid zones (Sánchez-Guillén et al., 2011, 2012; Swaegers, Sánchez-Guillén, Chauhan, et al., 2022; Arce-Valdés, Ballén-Guapacha, et al., 2025). (iii) Evaluate the consistency of outlier loci across hybrid zones by comparing and functionally annotating markers that exhibit excessive gene flow from one parental species into the other. If genomic architecture imposes shared constraints on introgression, we expect loci showing excess introgression to be associated with similar functional categories across hybrid zones, even if the specific loci involved differ among regions (Gompert et al., 2012; Edelman et al., 2019; Taylor & Larson, 2019).

## MATERIAL AND METHODS

### Sampling design, RAD-seq data processing and SNP calling

We used genomic RAD-sequencing data of *I. elegans* and *I. graellsii* previously published by Swaegers (2021) which provide dense genome-wide coverage across allopatric populations and multiple hybrid zones differing in their evolutionary dynamics. These data comprise 26 populations sampled between 2005 and 2015, including the allopatric distribution of *I. graellsii* in Spain, Portugal, and the Maghreb in northern Africa (n = 46), six populations from the allopatric distribution of *I. elegans* ranging from France and the United Kingdom to central Europe (n = 47), five populations from the North-west (NW) hybrid zone (n = 49), seven populations from the North-central (NC) hybrid zone (n = 68), and four populations from the South-east (SE) hybrid zone (n = 27; Figure 1; Supplementary Table 1).

**Figure 1.**
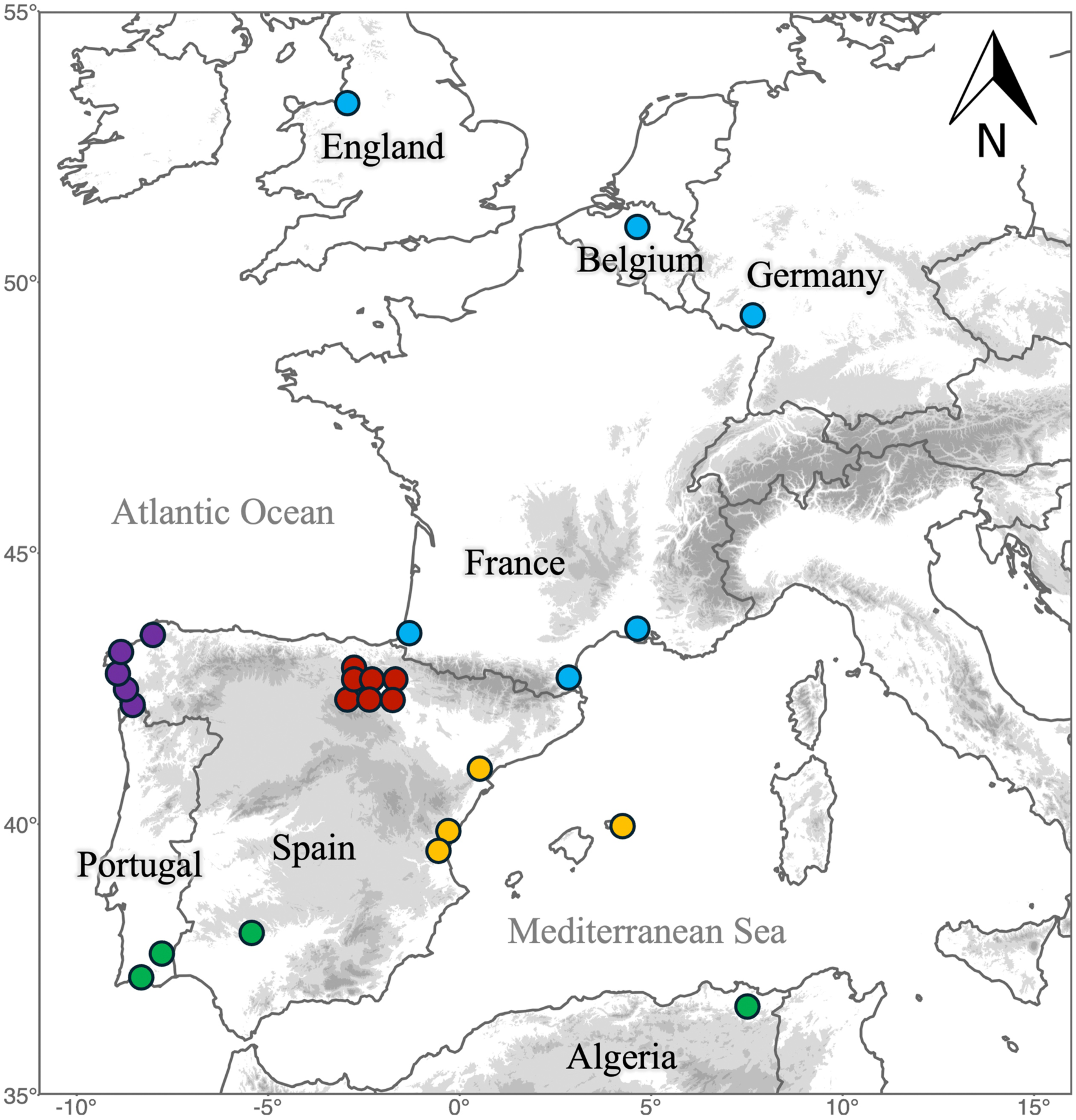
Geographic distribution of allopatric and hybrid sampling localities of *I. elegans* and *I. graellsii* across Europe and North Africa. Colours indicate population type: Green, allopatric *I. graellsii*; blue, allopatric *I. elegans*; yellow, South-east hybrid zone; purple, North-west hybrid zone; and red, North-central hybrid zone.

We assembled these sequences following the Stacks v2.2 pipeline (Catchen et al., 2013; Rochette et al., 2019). First, we demultiplexed and quality-filtered raw RAD-seq reads with *process_radtags*, removed PCR duplicates using *clone_filter* with default parameters, and aligned the cleaned reads to the most recent *I. elegans* reference genome (Price et al., 2022) using the BWA-MEM algorithm implemented in BWA v0.7.17 (Li & Durbin, 2009). We processed the resulting BAM files with *gstacks* and called variants using *populations* with the *–ordered-export* flag, filtering variants to retain a minimum minor allele frequency of 0.02. Using VCFtools v0.1.16 (Danecek et al., 2011), we further filtered the resulting VCF file to include only bi-allelic SNPs, recovering a total of 2,177,590 markers.

Following O’Leary et al. (2018), we tested several filtering schemes to retain an optimal number of high-quality SNPs and samples (Supplementary File 1). We selected the scheme retaining genotypes with a minimum sequencing depth of 5, loci with a maximum of 10% missing data, and samples with a maximum of 15% missing data, which resulted in 10,093 SNPs. We pruned markers for linkage disequilibrium using Plink v1.90 (Purcell et al., 2007) with the *--indep-pairwise* function, applying a sliding window of 50 SNPs and a step size of 5 SNPs. Pairwise linkage disequilibrium was estimated among SNPs within each window, and one random SNP was retained when the genotypic correlation coefficient (r²) exceeded 0.5. We then used allopatric samples as references for a preliminary supervised Admixture v1.3.0 analysis (Alexander & Lange, 2011) and classified individuals from sympatric populations as putative pure-species samples when their Q values were ≥ 0.999 for either parental species. We tested Hardy–Weinberg disequilibrium across the full distribution of both species using these pure-species samples in VCFtools and removed SNPs that showed significant deviation from Hardy–Weinberg equilibrium in both species simultaneously (p < 0.05). Finally, for subsequent analyses, we divided the full SNP dataset into autosomal (6,922 SNPs) and X-linked (236 SNPs) subsets, both comprising 237 samples.

### Demographic history inference

To investigate the origins of the three hybrid zones and reconstruct the demographic history of both species in Spain, we tested alternative scenarios of secondary contact and range expansion that could account for the contrasting evolutionary outcomes observed among hybrid zones. We employed the software DIYABC-RF (Collin et al., 2021) following best-practice recommendations for demographic inference using genomic data (Marchi et al., 2021). This framework combines Approximate Bayesian Computation with a Random Forest algorithm to compare observed genetic data against simulated datasets, enabling the identification of the most likely demographic scenarios. In addition, the use of machine learning enhances both the accuracy and robustness of model selection and parameter estimation, particularly in complex evolutionary histories involving divergence, gene flow, and admixture (Collin et al., 2021). This integrative approach is especially well-suited to disentangle the demographic processes underlying hybrid zone formations in systems characterised by recent divergence, ongoing gene flow, and spatially structured secondary contact (Csilléry et al., 2010; Collin et al., 2021), as in *I. elegans* and *I. graellsii*.

Using the autosomal dataset (6,922 SNPs), we compared eight alternative demographic scenarios that differed mainly in whether the hybrid zones had a single origin or originated independently or alternatively resulted from sequential colonisation events involving already established hybrid zones. These scenarios would reflect alternative hypotheses about the temporal and spatial dynamics of secondary contact following the recent southward expansion of *I. elegans* (details in Supplementary Figure 1).

For each scenario, we performed 3,000 coalescent simulations and evaluated the fit between simulated and observed data using a principal component analysis of all summary statistics included in the software. We selected the best scenario via random forest classification using 1,000 trees, which allowed for accurate discrimination among competing models despite the high dimensionality of the data. Then, we did another 40,000 coalescent simulations under the best-supported scenario to estimate key demographic parameters, including timing of the cessation of gene flow between species (assuming 0.67 generations per year, i.e., 1.5 years per generation, according to Corbet et al. 2006), effective population sizes, and admixture rates. To determine whether this number of simulations was adequate, we implemented a random forest regression approach to estimate error metrics, comparing results from the 40,000 simulations with those from a randomly selected subset of 12,000 simulations. As no meaningful differences in error metrics were detected between the two simulation datasets, we considered that the results obtained with 40,000 simulations, using a random forest analysis composed of 1,000 trees and 30% of these simulations as out-of-bag testing samples, were sufficiently robust.

### Genomic cline analyses

To describe patterns of introgression between multiple loci in each hybrid zone, we performed genomic clines analyses based on genotype frequencies along a genome-wide admixture gradient. This approach allows the identification of loci showing excessive or restricted gene flow, thereby linking zone-specific demographic histories with heterogeneity in genomic introgression (Gompert & Buerkle, 2011). Genomic cline analyses were performed for both autosomal (6,922 SNPs) and X-linked (236 SNPs) datasets. However, the SE hybrid zone was excluded from these analyses since only introgressed *I. elegans* individuals, as identified from SNP genomic data, were included from this region (Arce-Valdés, Swaegers, et al., 2025), precluding the estimation of genomic clines based on both parental and admixed individuals. Before the analyses, we applied additional filtering criteria tailored to genomic cline analyses. We filtered markers using the allopatric parental populations of *I. elegans* and *I. graellsii* as references, retaining only SNPs with less than 20% missing data in each parental group, an allele frequency difference greater than 0.10 between species and a minor allele frequency above 0.20. These criteria were applied to enrich the dataset for loci showing contrasting allele frequencies between the allopatric reference groups. Genomic clines were then modelled using the R package gghybrid (Bailey, 2022).

First, we estimated hybrid indices using a single Markov chain Monte Carlo (MCMC) run with 3,000 iterations and a burn-in of 1,000 iterations. Hybrid indices were then used to fit genomic clines through a second MCMC with 5,000 iterations and a burn-in of 2,000 iterations. In both analyses, MCMC stability was assessed using real-time trace plots. Finally, statistically significant differences in cline steepness and centre between NW and NC hybrid zones were assessed using a paired Wilcoxon signed-rank test (Wilcoxon, 1945) in both autosomes and X-linked loci. On the other hand, statistically significant differences in cline steepness between autosomes and X-linked loci in NW and NC hybrid zones were assessed using a Wilcoxon rank-sum test (Wilcoxon, 1945).

### Functional annotation of introgressed loci

To explore the heterogeneity of introgression across the genome and assess whether similar genomic regions are involved across hybrid zones, we classified outlier SNPs based on their cline parameters (Bailey, 2022). Specifically, SNPs with cline steepness values lower than one were classified as showing excessive gene flow, whereas SNPs with higher values were classified as showing restricted introgression. Additionally, based on cline centre estimates, SNPs with centre values greater than 0.5 were interpreted as carrying *I. graellsii* alleles in an *I. elegans* genomic background, whereas values lower than 0.5 indicated *I. elegans* alleles in an *I. graellsii* background. Loci were considered outliers only when these patterns were statistically significant (p < 0.05; Bailey, 2022). We tested for statistically significant differences in the frequency of these outlier classes using Fisher’s exact test (Fisher, 1922). Finally, we functionally annotated outlier SNPs showing excessive introgression to evaluate whether shared or zone-specific introgression patterns involved genes associated with similar biological functions. We mapped outlier SNPs to the GFF annotation file of the *I. elegans* reference genome (Price et al., 2022) and identified annotated coding sequence (CDS), mRNA, or exon features that directly contained the genomic position of each outlier SNP. Given the reduced representation nature of RAD-seq data, only the SNP positions themselves were considered, and surrounding genomic regions were not evaluated. For each feature, we retrieved the corresponding protein sequences from GenBank using NCBI E-utilities (Sayers, 2022). When outlier SNPs were located within features associated with multiple homologs, isoforms, or protein variants, we randomly selected a single representative protein sequence per SNP. Protein sequences were then analysed using InterProScan v.5 (Jones et al., 2014) to identify conserved protein domains based on integrated protein signature databases. Gene Ontology (GO) terms were assigned to each sequence according to the functional annotations associated with the detected InterPro entries.

## RESULTS

### Demographic history inference

We compared eight alternative demographic scenarios describing the formation of the SE, NW, and NC hybrid zones, considering either independent admixture events or sequential formation mediated by migration among hybrid zones (Supplementary Figure 1). Using 3,000 simulations per scenario, all proposed models were consistent with the observed dataset based on PCA analysis of summary statistics (Supplementary Figure 2). Among the tested models, Scenario 1, describing the hybrid zones as independently formed admixture events, with the SE hybrid zone forming first, followed by the NW, and most recently the NC hybrid zone, received the highest support according to random forest votes (517 votes, posterior probability = 0.456; Figure 2; Table 1). Under this scenario, the SE hybrid zone showed a low genetic contribution from *I. graellsii* (4.5%). Admixture proportions were more balanced in the NW (*I. graellsii* = 54.3%, *I. elegans* = 45.7%), and in the NC (*I. graellsii* = 53.5%, *I. elegans* = 46.5%; Figure 2).

**Figure 2.**
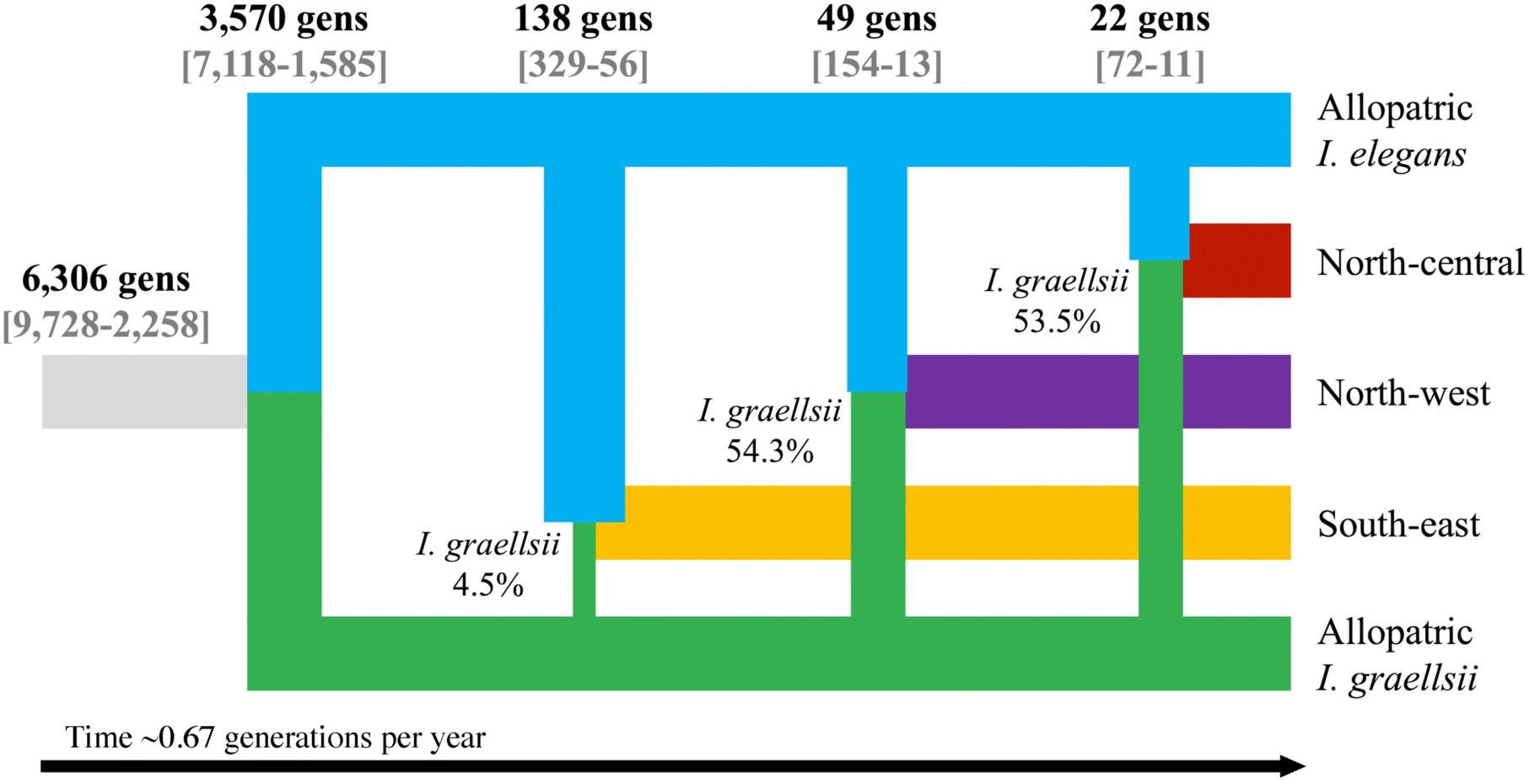
Best-supported demographic scenario inferred using DIYABC-RF for the formation of the three hybrid zones. Time is shown in generations before present, with the earliest time point representing a demographic expansion preceding the cessation of gene flow between lineages. The model supports independent origins for each hybrid zone, with the South-east hybrid zone forming first, followed by the North-west hybrid zone and then the North-central hybrid zone. Admixture proportions represent the genetic contribution of *I. graellsii* to each hybrid zone, with the remaining proportion corresponding to *I. elegans*.

**Table 1.**
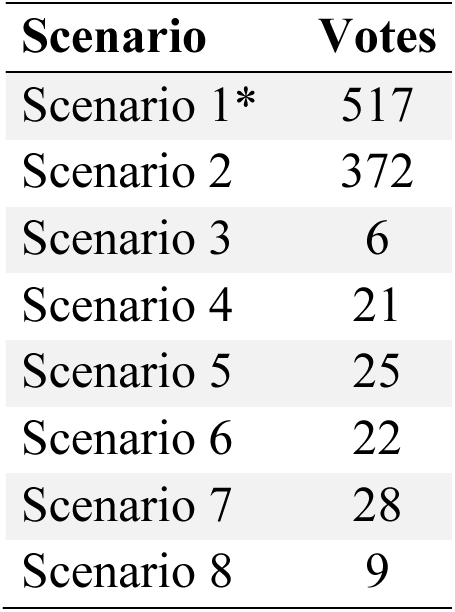
Random forest vote counts for the eight demographic scenarios tested in the DIYABC-RF analysis. Scenario 1 received the highest number of votes, indicating the strongest support among all models based on the simulated datasets.

| Scenario | Votes |
| --- | --- |
| Scenario 1* | 517 |
| Scenario 2 | 372 |
| Scenario 3 | 6 |
| Scenario 4 | 21 |
| Scenario 5 | 25 |
| Scenario 6 | 22 |
| Scenario 7 | 28 |
| Scenario 8 | 9 |

A demographic expansion preceding the cessation of gene flow was inferred around 6,306 generations ago (90% CI: 9,728–2,258), corresponding to around 9,459 years ago (90% CI: 14,592–3,387; assuming 0.67 generations per year, i.e., 1.5 years per generation, Corbet et al., 2006), during this period, effective population size increased from 881 at the onset of the expansion to present-day allopatric population sizes estimated for *I. elegans* (6,130) and *I. graellsii* (6,904; Figure 2; Table 2). The cessation of gene flow between *I. elegans* and *I. graellsii* subsequently estimated at approximately 3,570 generations ago (90% CI: 7,118–1,585; Figure 2; Table 2), corresponding to 5,355 years ago (90% CI: 10,677–2,378). More recently, parameter estimates indicate that the SE hybrid zone was established approximately 138 generations ago (90% CI: 329–56; Figure 2; Table 2), corresponding to 207 years ago (90% CI: 494–84). The NW hybrid zone formed more recently, around 49 generations ago (90% CI: 154–13; Figure 2; Table 2), equivalent to 73.5 years ago (90% CI: 231–19.5), followed by the NC hybrid zone, which formed approximately 22 generations ago (90% CI: 72–11; Figure 2; Table 2), corresponding to 33 years ago (90% CI: 108–16.5). Finally, effective population sizes estimated for the SE (Ne = 6,014) and NW (Ne = 6,592) hybrid zones were similar, whereas the NC showed a smaller effective population size (Ne = 4,048; Table 2).

**Table 2.** Predicted parameter estimates from the DIYABC-RF analysis under the selected demographic scenario 1. Parameters include t: times of species divergence and hybrid zone formation, Ne: effective population sizes, r: admixture proportions representing the genetic contribution of *I. graellsii* (with the complement corresponding to *I. elegans*).

| Parameter | Median | Lower CI | Upper CI |
| --- | --- | --- | --- |
| t ancestral | 6,306 | 2,258 | 9,728 |
| t divergence | 3,570 | 1,585 | 7,118 |
| t South-east | 138 | 56 | 329 |
| t North-west | 49 | 13 | 154 |
| t North-central | 22 | 11 | 72 |
| Ne ancestral | 881 | 126 | 9,415 |
| Ne allopatric *I. elegans* | 6,130 | 2,258 | 9,573 |
| Ne allopatric *I. graellsii* | 6,904 | 3,070 | 9,683 |
| Ne South-east | 6,014 | 1,830 | 9,597 |
| Ne North-west | 6,592 | 2,236 | 9,319 |
| Ne North-central | 4,048 | 459 | 9,183 |
| r South-east | 0.045 | 0.013 | 0.106 |
| r North-west | 0.543 | 0.414 | 0.688 |
| r North-central | 0.535 | 0.398 | 0.673 |

### Genomic cline analyses

#### Autosomal genomic clines

For the autosomal SNP dataset, genomic cline analyses revealed no significant differences in cline steepness between the NW (median = 2.636) and NC (median = 2.64; Figure 3a) hybrid zones (paired Wilcoxon signed-rank test: V = 882,637; p = 0.379; Figure 3a; Supplementary Figure 3). In contrast, the genomic cline centres differed significantly between hybrid zones (paired Wilcoxon signed-rank test: V = 712,398; p < 0.001). The median cline centre was 0.48 in the NW hybrid zone and 0.59 in the NC hybrid zone (Figure 3b).

**Figure 3.**
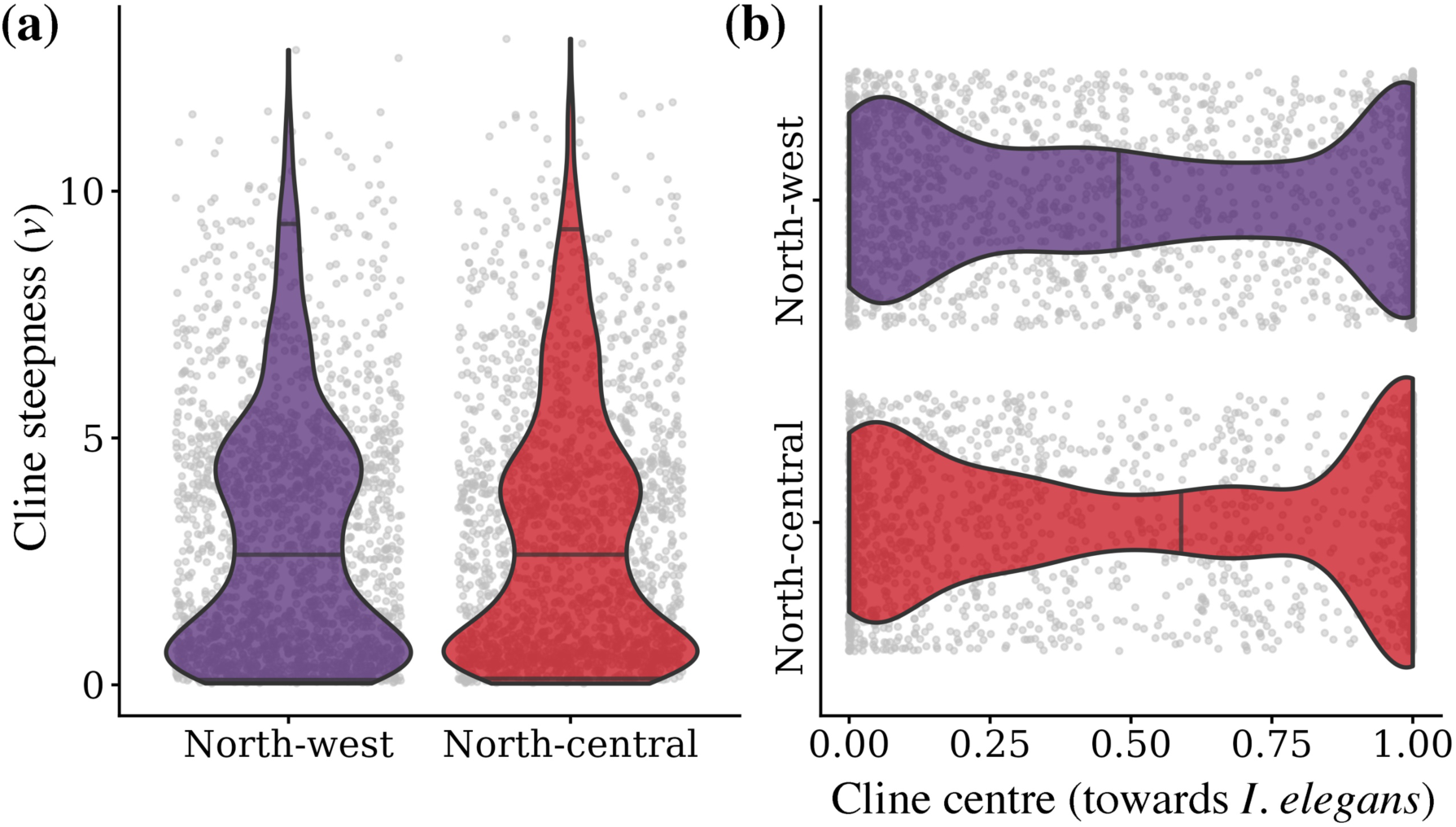
Genomic cline variation across hybrid zones in *I. elegans* and *I. graellsii*. Violin plots of cline steepness (a) and cline centre (b) for the North-west and North-central hybrid zones based on the autosomal SNP dataset. Lines within each violin indicate the median and the 0.025 and 0.975 quantiles, and individual SNPs are shown in grey dots. Cline steepness reflects the strength of deviation from genome-wide neutral expectations of introgression, with higher values indicating steeper transitions across the admixture gradient. Cline centre represents the position of allele frequency change along the hybrid index, indicating the direction of introgression between parental genomic backgrounds.

Outlier analyses identified differences in the number of autosomal loci with excessive introgression patterns between hybrid zones. A similar number of autosomal SNPs showing excessive gene flow were detected in the NC (n = 48) compared to the NW hybrid zone (n = 42; Supplementary Figure 4a). Of these, four outlier SNPs were shared between both hybrid zones. Conversely, four SNPs associated with restricted gene flow were detected in NW (Supplementary Figure 3a), and one in the NC hybrid zone (Supplementary Figure 3b).

Directional introgression analyses identified asymmetries between hybrid zones. We detected 133 SNPs originating from *I. elegans* introgressed into the genome of *I. graellsii* in the NW and 154 in the NC hybrid zone, with 43 SNPs shared between both hybrid zones (Supplementary Figure 4b). Notably, a substantially higher number of SNPs showed introgression in the opposite direction. Specifically, 304 SNPs from *I. graellsii* were detected in the genome of *I. elegans* in NW and 442 in NC hybrid zone, of which 163 were shared between the two hybrid zones (Supplementary Figure 4c).

After merging both types of outliers, including SNPs showing excessive gene flow and those exhibiting bidirectional or unidirectional introgression, we detected a greater number of SNPs with excessive bidirectional introgression in the NC and NW hybrid zones (20 unique to NC, 27 unique to NW, and two shared; Figure 4a). Similarly, numerous SNPs showed unidirectional introgression from *I. graellsii* into *I. elegans* (21 unique to NC, 8 unique to NW, and one shared; Figure 4b). In contrast, far fewer SNPs showed introgression in the opposite direction, from *I. elegans* into *I. graellsii* (three unique to NC, two unique to NW, and one shared; Figure 4c).

**Figure 4.**
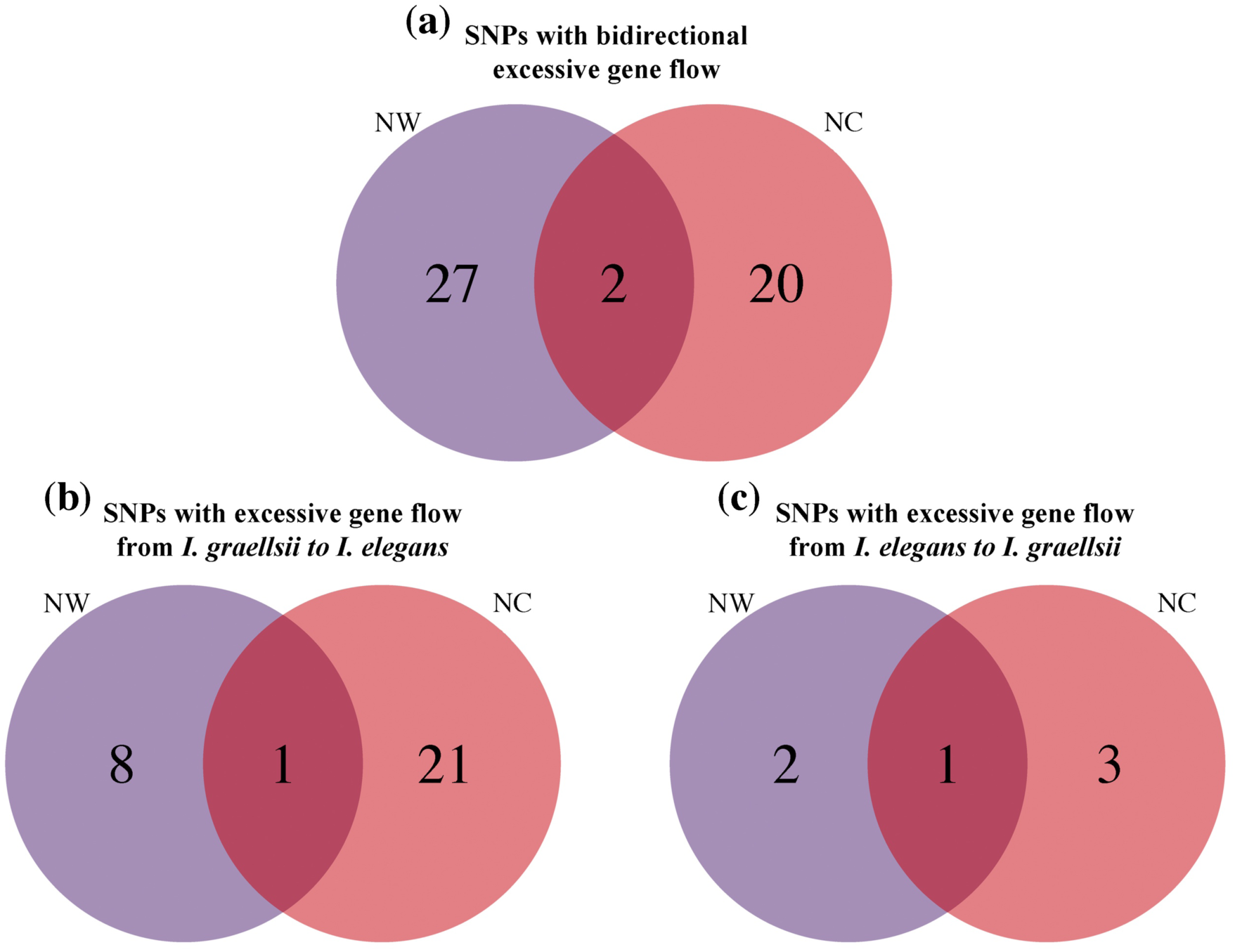
Asymmetric genomic introgression between *I. elegans* and *I. graellsii*. Outlier SNPs showing excessive bidirectional introgression (a) and excessive directional introgression between parental species, with (b) introgression of *I. graellsii* alleles into the *I. elegans* genomic background, and (c) introgression of *I. elegans* alleles into the *I. graellsii* genomic background.

#### X-linked genomic clines

For the X-linked SNP dataset, genomic cline steepness (Supplementary Figure 5) differed between hybrid zones (paired Wilcoxon signed-rank test: V = 2,106; p = 0.03). Slopes were steeper in the NC (median = 4.57) than in the NW hybrid zone (median = 4.12; Supplementary Figure 6a). Furthermore, the genomic cline centre median was shifted toward *I. elegans* in NC (median = 0.67) and toward *I. graellsii* in NW hybrid zone (median = 0.40; Supplementary Figure 6b), with the difference between zones being statistically significant (paired Wilcoxon signed-rank test: V = 1,429; p = 0.003). Furthermore, no X-linked SNPs were identified as excessive gene flow in NW and only one in NC hybrid zone.

Finally, when comparing cline steepness between autosomal and X-linked SNPs, we found that X-linked loci exhibited significantly steeper clines than autosomal loci in both hybrid zones. In the NW hybrid zone, the median steepness was 2.63 for autosomal SNPs and 4.12 for X-linked SNPs (Wilcoxon rank-sum test: W = 74,482; p < 0.001). Similarly, in the NC hybrid zone, median steepness was 2.64 for autosomal loci and 4.57 for X-linked loci (Wilcoxon rank-sum test: W = 63,932; p < 0.001). This pattern suggests a stronger restriction to introgression on the sex chromosome relative to the autosomes.

### Functional annotation of introgressed loci

Among the 75 autosomal SNPs showing excess unidirectional or bidirectional introgression between parental species, we found 19 protein sequences associated with 87 Gene Ontology terms (Supplementary Table 2) representing several shared functional categories (Supplementary Figure 7). Terms associated with protein-protein interactions and catalytic activity were detected in both hybrid zones, as well as processes related to nucleotide metabolism and signalling pathways. Despite these functional similarities, some regional trends were evident. In the NW hybrid zone, annotations were more frequently associated with cellular structural organisation, including cytoskeletal and extracellular matrix-related processes, whereas in the NC hybrid zone, functions were more associated in metabolic and redox-related pathways.

At the molecular function level, both hybrid zones shared protein binding, protein–protein interactions and general catalytic activities as the most common annotations. The NW hybrid zone additionally showed a higher representation of signalling-related functions, including kinase activity and receptor-mediated processes, whereas the NC hybrid zone was associated with cofactor-binding functions and enzymes involved in metabolic pathways, as those associated with aminoacid metabolism and redox processes.

At the biological process level, both hybrid zones encompassed core cellular and regulatory functions. However, the NW hybrid zone was more strongly associated with processes related to cellular architecture and structural organisation, while the NC hybrid zone showed a greater contribution of metabolic processes and RNA-related regulation.

Finally, the cellular component category indicated that proteins associated with these outliers SNPs are broadly distributed across intracellular compartments in both hybrid zones, primarily within the nucleus and cytoplasm. Nevertheless, NW hybrid zone included more components related to the extracellular environment and specialised cellular structures, whereas the NC hybrid zone showed a broader representation of organelle-associated components. Overall, these annotations indicate that outlier-associated loci in both hybrid zones are associated to fundamental cellular functions, with zonal differences reflecting variation in the relative contribution of structural, regulatory, and metabolic processes.

## DISCUSSION

Why hybridisation leads to divergent outcomes remains a central question in evolutionary biology. Our results show that recent divergence, spatially replicated but temporally distinct secondary contact, and chromosome-specific permeability to gene flow jointly shape the demographic trajectories and introgression patterns observed across the three hybrid zones between *I. elegans* and *I. graellsii* in Spain. Consistent with our predictions, hybrid zones originated independently and at different times, genomic clines varied in direction and magnitude among zones, and loci showing excess introgression shared broad functional categories despite regional differences.

### Demographic history and genomic architecture jointly shape introgression patterns

Post-glacial climatic fluctuations likely shaped the evolutionary history of *I. elegans* and *I. graellsii*. Phylogenetic estimates place the divergence between these species within the last ∼100,000 years (Sánchez-Guillén et al., 2020; Blow et al., 2021), with our demographic analyses indicating a more recent cessation of gene flow between 2,378 and 10,677 years ago, suggesting a prolonged history of interaction following initial divergence within the Holocene. This timing coincides with climatic warming following the Last Glacial Maximum (∼21,000 years ago), which reshaped species distributions across Europe (Hewitt, 2000). Given that the Iberian Peninsula served as a significant glacial refugium (Gómez & Lunt, 2007; Hewitt, 1999), the common ancestor of both species may have persisted there, with subsequent post-glacial range expansion promoting the differentiation between the Iberian/North Africa restricted *I. graellsii* and the widely distributed *I. elegans*. The strong dispersal ability and ecological flexibility of *I. elegans* likely facilitated rapid northward expansion and local adaptation across contrasting environments, ranging from the warm and human-modified agricultural ponds in the Iberian Peninsula (Swaegers, Sánchez-Guillén, Carbonell, et al., 2022; Wellenreuther et al., 2018) to cold freshwater systems in northern Europe (Dudaniec et al., 2018; Lancaster et al., 2015). Such post-glacial divergence coupled with rapid range expansion is increasingly recognised as a recurrent pattern in Odonata, including *Ischnura*, *Calopteryx*, and *Enallagma*, where high dispersal ability and ecological flexibility promote shallow genetic divergences, demographic expansions, and secondary contact among closely related lineages (Turgeon & McPeek, 2002; Wellenreuther & Sánchez-Guillén, 2016). Similar post-glacial divergence and expansion scenarios have been documented across a wide range of taxa such as insects (Capblancq et al., 2015, 2019; Mas-Peinado et al., 2022; Schmitt et al., 2006), fishes (Schluter & McPhail, 1992; Volckaert et al., 2002), and birds (Ellegren et al., 2012; Sætre & Sæther, 2010; Warmuth et al., 2021), indicating that recent divergence followed by secondary contact is a recurrent outcome of post-glacial range dynamics.

Demographic modelling indicates that secondary contact occurred independently across hybrid zones under distinct temporal and demographic contexts, providing a framework to interpret heterogeneous introgression patterns. DIYABC-RF estimates suggest the earliest contact occurred in the SE hybrid zone ∼207 years ago, consistent with the first records of *I. elegans* in Spain (Ocharan-Larrondo, 1987) and with large effective population sizes and long-term asymmetric introgression (Sánchez-Guillén et al., 2011). In contrast, the NW (∼74 years ago) and NC (∼33 years ago) zones represent more recent contacts with initially balanced admixture but divergent introgression trajectories between them —bidirectional in NW and unidirectional in NC (Arce-Valdés, Swaegers, et al., 2025; Swaegers, Sánchez-Guillén, Chauhan, et al., 2022). Together, these temporally structured histories support a mosaic hybrid zone framework (Rand & Harrison, 1989), where variation in the timing and demographic context of secondary contact shapes genomic heterogeneity, introgression patterns, and the evolution of reproductive isolation across zones (Arce-Valdés, Ballén-Guapacha, et al., 2025; Ballén-Guapacha et al., 2024).

At the same time, our results indicate that genomic architecture strongly influences where gene flow is constrained or facilitated. Autosomal loci show widespread excess introgression, but only a small fraction of these loci is shared among hybrid zones, suggesting that autosomal gene flow is context dependent and shaped by local demographic and ecological conditions rather than intrinsic species differences. In contrast, X-linked loci showed reduced permeability to introgression compared to autosomes and follow neutral genomic cline expectations. This neutral pattern of the X chromosome, which has been previously reported in this system (Swaegers, Sánchez-Guillén, Chauhan, et al., 2022), supports the view that sex chromosomes often act as stable barriers to gene flow (Horta et al., 2025; Presgraves, 2018), although demographic history may further modulate their resistance to introgression.

Beyond these genome-wide introgression patterns, which are consistent with previous studies in this hybrid system that documented contrasting hybridisation dynamics and evolutionary outcomes between the NC and NW hybrid zones (Arce-Valdés, Ballén-Guapacha, et al., 2025; Arce-Valdés, Swaegers, et al., 2025; Ballén-Guapacha et al., 2024). This aligns our results with a growing body of literature showing that genomic permeability to introgression can vary geographically within the same species pair (Harrison & Larson, 2014; Taylor et al., 2015). This variation can be driven by factors such as effective population size, initial proportions of parental species, the strength of reproductive barriers at the onset of secondary contact, and the time since initial hybridisation (Barton & Hewitt, 1985; Payseur & Rieseberg, 2016). Consequently, similar barriers to gene flow can yield markedly different genomic outcomes across geographically replicated hybrid zones (Ravinet et al., 2017). From a broader perspective, these findings reinforce the view that hybrid zones are dynamic systems that change through time rather than static endpoints of species interactions. The recent divergence of these species—likely associated with post-glacial processes—and their subsequent secondary contact and rapid range expansion in the Iberian Peninsula under contemporary climate warming (Arce-Valdés, Swaegers, et al., 2025) make this system particularly informative for understanding speciation with gene flow under natural conditions. In line with patterns documented in other hybrid model systems—such as *Heliconius* butterflies (Edelman et al., 2019; Martin et al., 2013), *Bombina* toads (Fijarczyk et al., 2011), *Ficedula* flycatchers (Ellegren et al., 2012), and house mice (Macholán et al., 2011)—the contrasting genomic patterns observed among hybrid zones of different ages in *I. elegans* and *I. graellsii* illustrate how demographic history and genomic architecture jointly interact to shape introgression dynamics and speciation trajectories in evolving landscapes.

### Functional biases in genomic introgression

Patterns of introgression observed here are consistent with a growing body of work showing that gene flow across species boundaries is not random, but instead biased toward particular functional classes of genes (Burgarella et al., 2019; Janoušek et al., 2015). Across a wide range of insect hybrid systems, loci involved in broadly acting cellular and regulatory functions —such as signalling, protein binding, transport, and general metabolic processes—are more likely to introgress than genes directly associated with reproductive isolation, which tend to remain resistant to gene flow (Horta et al., 2025), suggesting that repeatability in introgression often emerges at the level of gene function rather than gene identity.

Importantly, however, such conclusions have largely been drawn from interspecific comparisons, either across different species pairs or among independent hybrid systems (Edelman et al., 2019; Fontaine et al., 2015; Gompert et al., 2012; Martin et al., 2013; Taylor & Larson, 2019). In contrast, our results show that similar patterns of functional bias in introgression can also be detected at an intraspecific scale, across multiple hybrid zones formed between the same pair of species. Despite marked differences among hybrid zones in demographic history, directionality of gene flow, and the strength of reproductive barriers, loci showing excess introgression consistently belonged to similar functions. Notably, overlap among introgressed SNPs across hybrid zones was limited, indicating that repeatability emerges at the level of gene function rather than the repeated introgression of the same loci. Although some functional categories associated with introgressed loci were specific to individual hybrid zones, these patterns may reflect local adaptation of *I. elegans* to warmer colonised environments. Importantly, this suggests that even within a single species pair, convergence in introgression patterns can arise from functional similarity rather than shared genetic identity. Nevertheless, these results should be interpreted cautiously, and future analyses with datasets with higher genomic representation (e.g. Whole Genome Sequencing) will be necessary to robustly evaluate these patterns through gene ontology enrichment approaches; and to confirm whether the observed functional associations truly reflect local adaptive processes.

Finally, the functional categories associated with introgressed loci in our study also overlap with gene functions previously implicated in ecological and physiological responses in *I. elegans*. Previous work has shown that genes involved in regulation, metabolism, transport, and cellular signalling play a central role in mediating responses to thermal gradients, environmental stress, and rapid range expansion in this species (Dudaniec et al., 2018; Lancaster et al., 2015). In this context, the introgression of loci associated with broadly acting cellular functions is consistent with existing evidence that responses to new environments in *I. elegans* are underpinned by genes with general physiological roles, rather than narrowly specialised pathways.

### Conclusions

Our results show that hybridisation between *I. elegans* and *I. graellsii* reflects a dynamic interplay between demographic history and genomic architecture, generating heterogeneous introgression landscapes across independently formed hybrid zones. Variation in the timing of secondary contact, admixture proportions, and effective population sizes was associated with strongly zone-specific introgression dynamics, whereas chromosome architecture imposed consistent constraints: autosomes remained comparatively permeable to gene flow, while the X chromosome acted as a stable barrier. Notably, autosomal loci showing excess introgression differed among hybrid zones, yet converged on similar functional categories, revealing functional repeatability despite genomic contingency. Together, these findings suggest that demographic context drives geographic variation in introgression outcomes, while intrinsic genomic architecture and functional constraints can promote partly predictable evolutionary responses during early stages of speciation with gene flow.

## Data availability

All scripts used in this study are available in the OSF repository https://osf.io/hmz7k/overview?view_only=931486f31f0b4b69a025bf2407aa45da and raw genomic data are available in the Dryad repository https://doi.org/10.5061/DRYAD.GQNK98SP8.

## Author contributions

MS-P and LRA-V contributed to conceptualisation, data curation, formal analysis, investigation, methodology, software, validation, visualisation, and writing (original draft and review & editing). JEO-M contributed to investigation, methodology, validation, and writing (review & editing). JS, JRC-R, CG-R, EI-L, and BH contributed to validation and writing (review & editing). FB-D contributed to supervision, validation, and writing (review & editing). Additionally, EI-L and F-BD contributed to methodology. RAS-G contributed to conceptualisation, investigation, resources, supervision, validation, and writing (original draft and review & editing). Finally, MS-P and RAS-G contributed to project administration.

## Acknowledgments

We thank Janet Nolasco Soto and Emanuel Villafán de la Torre for technical support. Bioinformatic analyses were performed on the Huitzilin 2.0 HPC system at the Instituto de Ecología A.C. (INECOL). This research forms part of MS-P’s PhD thesis.

---

**Supplementary Figure 1.**
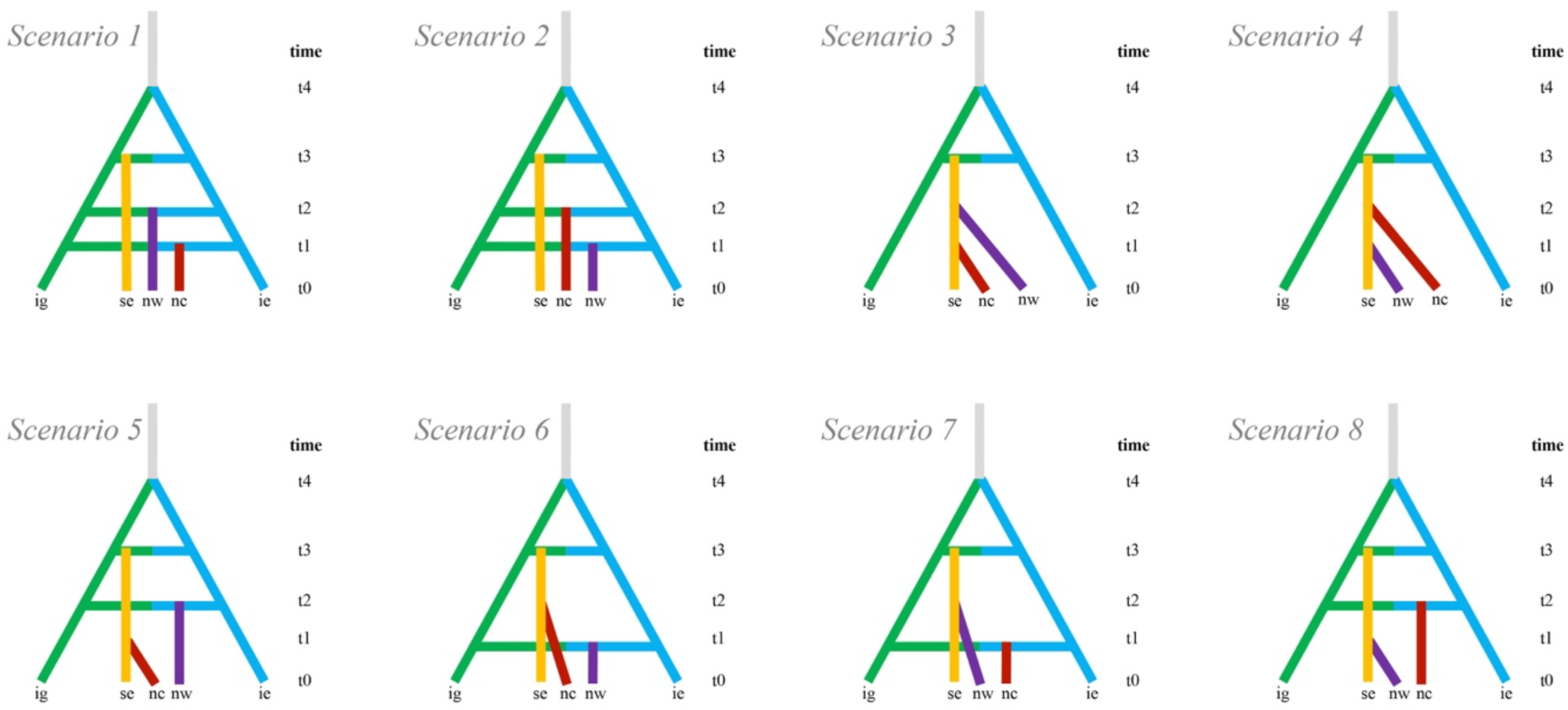
Schematic overview of the eight alternative demographic scenarios tested using DIYABC-RF. Each scenario represents a different hypothesis about the timing and origin of the three hybrid zones, considering either independent admixture events or sequential formation driven by migration. Scenarios 1–2 represent independent origins of the NW, NC and SE hybrid zones, differing in the order of divergence. Scenarios 4–8 represent non-independent origins, where one zone arise from the split of other after divergence. In all scenarios, time points (t0–t4) indicate divergence or admixture events to the establishment of each hybrid zone. ig: *I. graellsii*, ie*: I. elegans*, nw: North-west hybrid zone, nc: North-central hybrid zone, se: South-east hybrid zone.

**Supplementary Figure 2.**
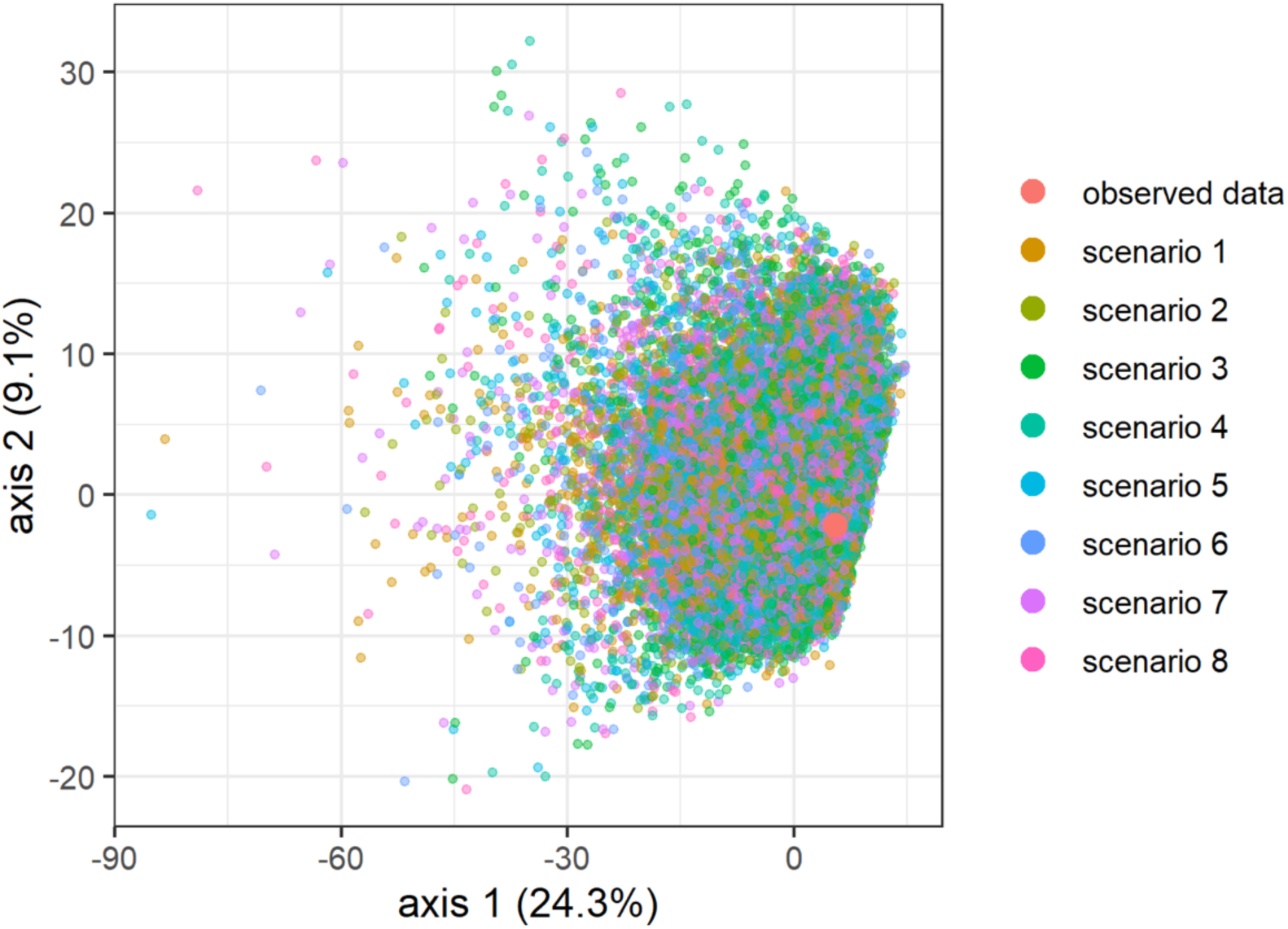
Principal component analysis plot for model checking of the DIYABC-RF analysis derived from the simulated training datasets across the eight demographic scenarios. The observed dataset (red circle) falls well within the cloud of simulated points, indicating that the observed summary statistics are consistent with the prior parameter distributions and model space explored.

**Supplementary Figure 3.**
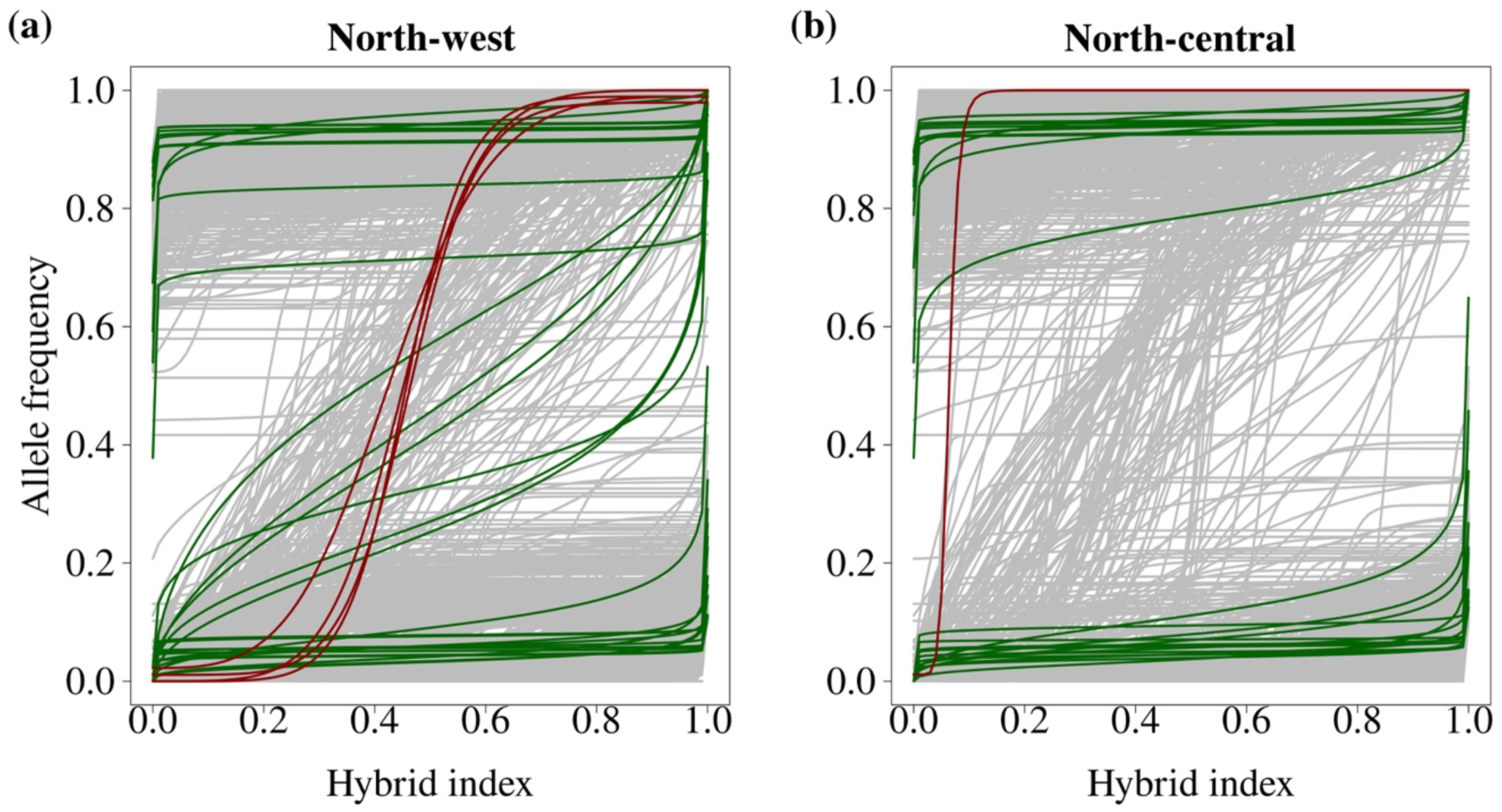
Genomic clines estimated for autosomal SNP dataset across (a) NW and (b) NC hybrid zone. Clines are coloured according to their inferred introgression category, with loci showing neutral introgression rates (grey), excessive introgression (green), and restricted gene flow (red).

**Supplementary Figure 4.**
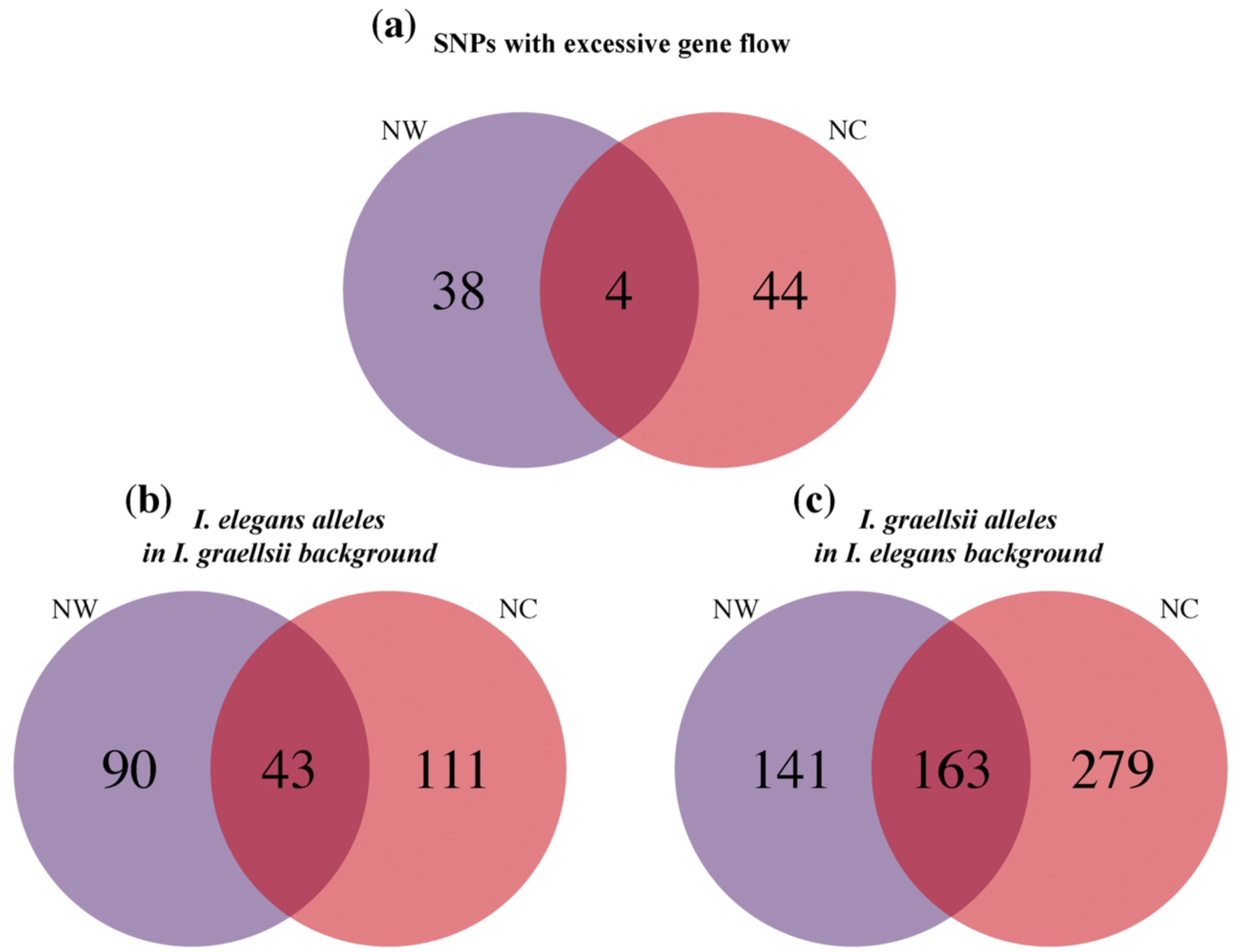
Comparison of the autosomal outlier SNPs in NW (purple circle) and NC (red circle) hybrid zones. (a) Cline steepness outlier SNPs showing excessive introgression; (b) cline centre outlier SNPs from *I. elegans* into the genomic background of *I. graellsii*; and (c) *vice versa*.

**Supplementary Figure 5.**
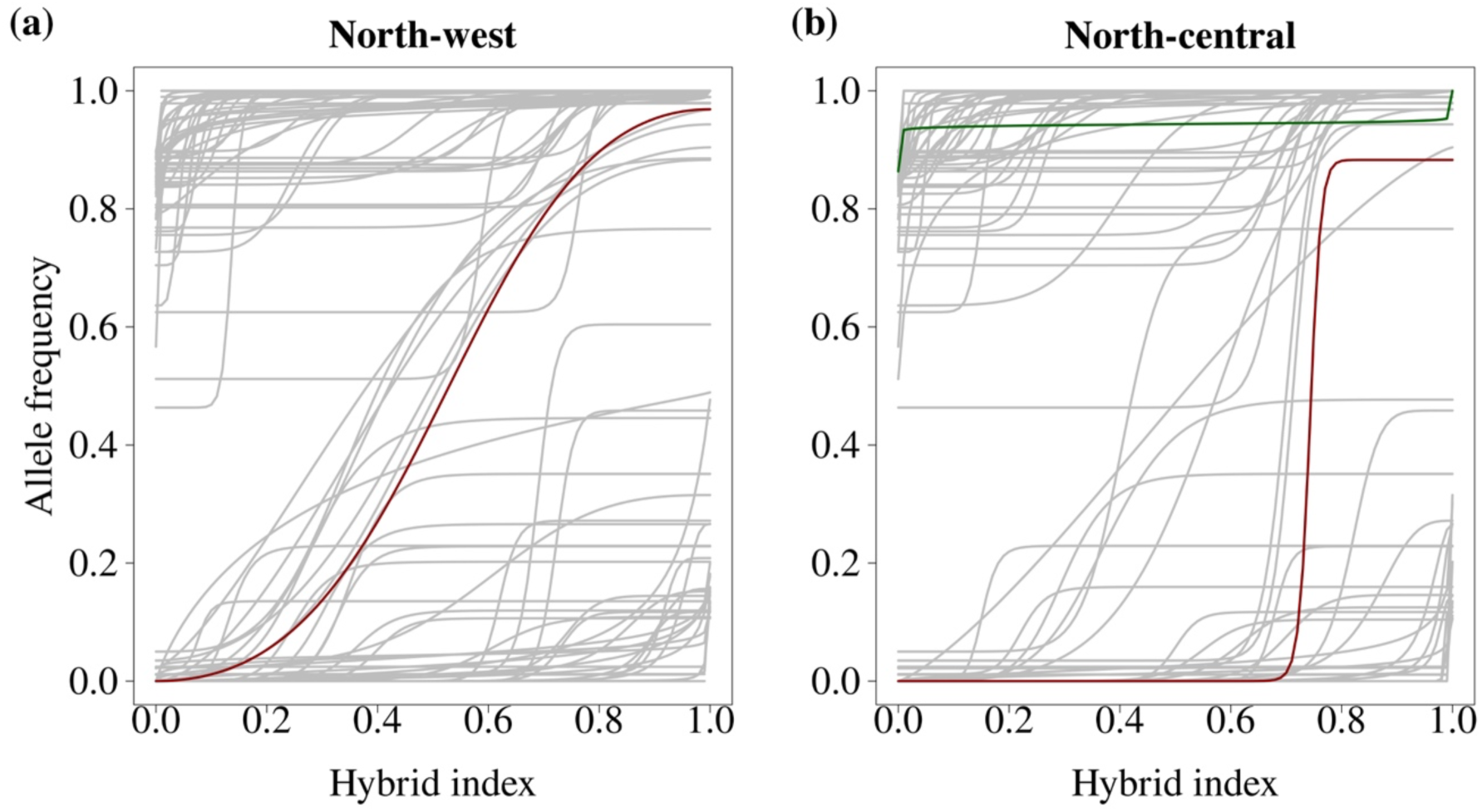
Genomic clines estimated for X-linked SNP dataset across (a) NW and (b) NC hybrid zone. Clines are coloured according to their inferred introgression category, with loci showing neutral introgression rates (grey), excessive introgression (green), and restricted gene flow (red).

**Supplementary Figure 6.**
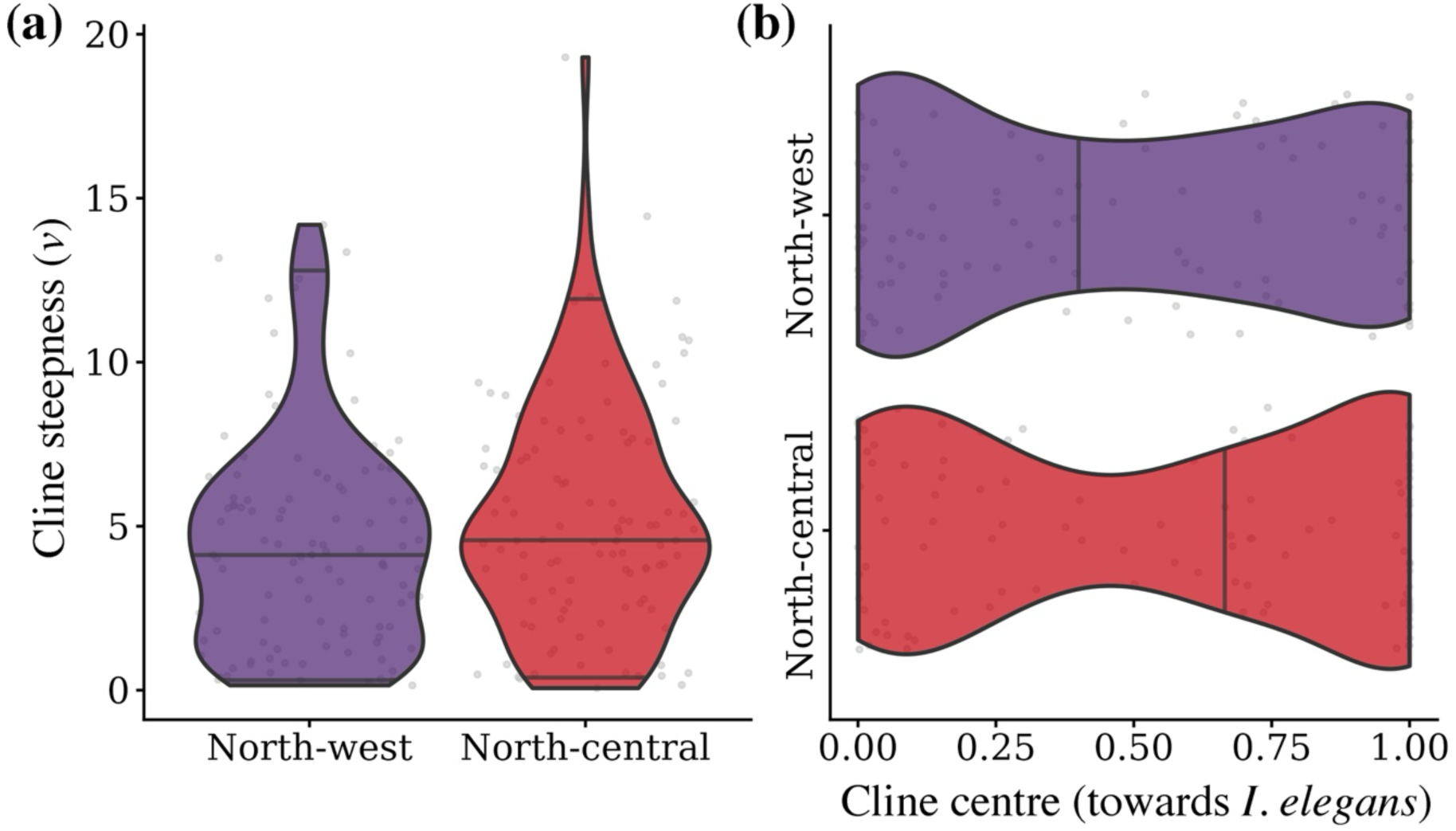
Violin plots of cline steepness (a) and cline centre (b) for the North-west and North-central hybrid zones for the X-linked SNPs dataset. Lines within each violin indicate the median and the 0.025 and 0.975 quantiles, and individual SNPs are shown in grey dots. Cline steepness reflects the strength of deviation from genome-wide neutral expectations of introgression, with higher values indicating steeper transitions across the admixture gradient. Cline centre represents the position of allele frequency change along the hybrid index, indicating the direction of introgression between parental genomic backgrounds.

**Supplementary Figure 7.**
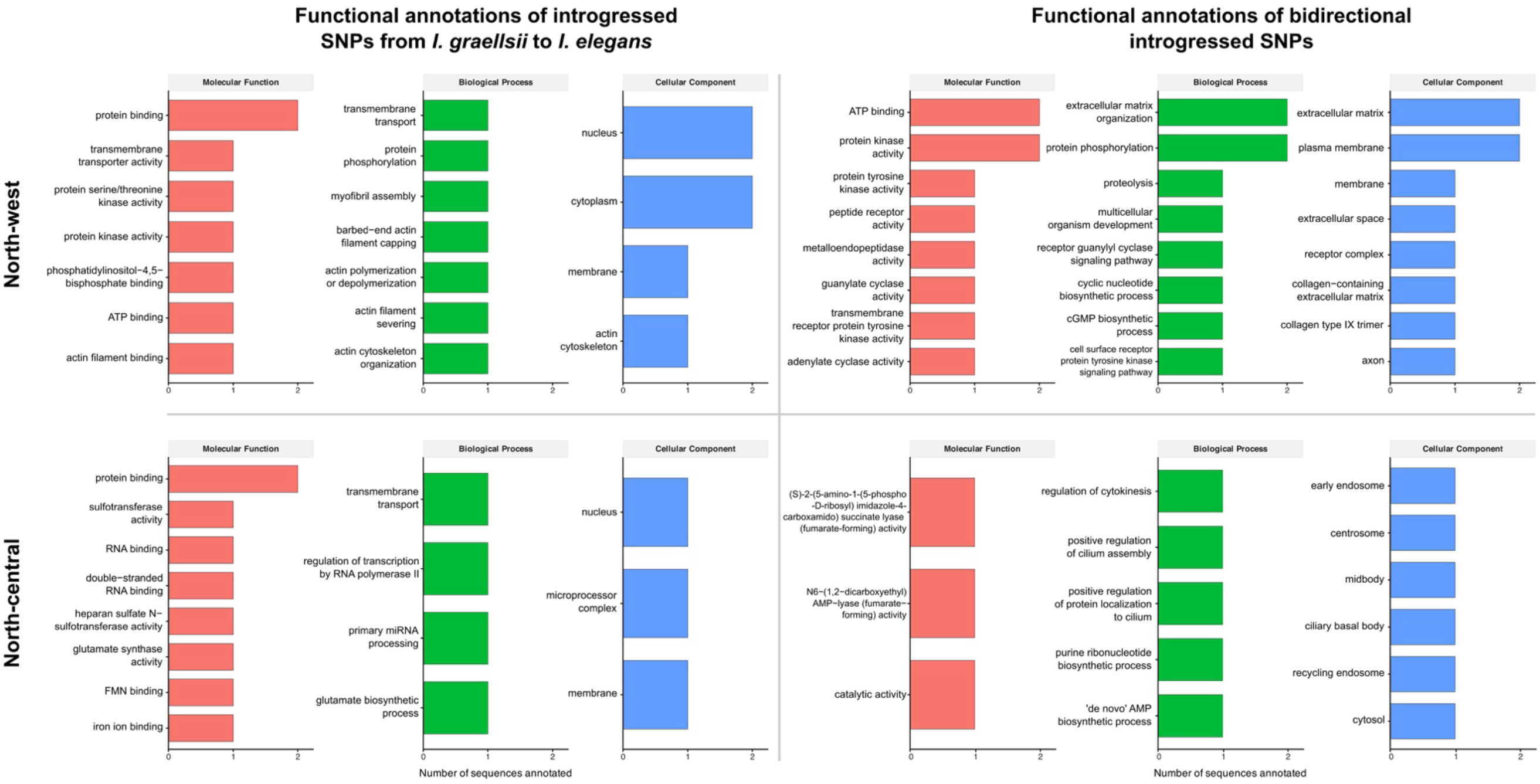
Frequency of Gene Ontology (GO) terms associated with SNPs showing excessive introgression, grouped by GO categories. These functions are shown for SNPs with excessive introgression from *I. graellsii* into *I. elegans* and for bidirectional introgression, across both the North-west and North-central hybrid zones.

**Supplementary Table 1.** Sampling information for all populations included in the analyses. The region, locality name, geographic coordinates, sampling years, and the number of individuals retained in the full SNPs datasets are provided.

| Region | Locality | Latitude | Longitude | Year | Full SNPs |
| --- | --- | --- | --- | --- | --- |
| **Allopatric**<br>***I. graellsii*** | Seyhouse, Algeria | 36°28'27.03"N | 07°22'27.03"E | 2011 | 7 |
|  | Ribeira de Cobres, Portugal | 37°39'49.25"N | 07°57'48.10"W | 2005 | 8 |
|  | Guadito, Spain | 37°49'10.44"N | 05°03'03.02"W | 2008 | 10 |
|  | Salgados, Portugal | 37°05'29.70"N | 08°19'44.70"W | 2015 | 21 |
| **North-west**<br>**hybrid zone** | Cachadas, Spain | 42°27'12.19"N | 08°50'55.37"W | 2015 | 9 |
|  | Doniños, Spain | 43°29'23.96"N | 08°18'54.01"W | 2014 | 10 |
|  | Laxe, Spain | 43°12'46.82"N | 08°59'38.35"W | 2014 | 10 |
|  | Louro, Spain | 42°45'28.08"N | 09°05'44.90"W | 2013 | 10 |
|  | Xuño, Spain | 42°32'28.90"N | 09°01'34.40"W | 2014 | 10 |
| **North-central**<br>**hybrid zone** | Arreo, Spain | 42°46'39.01"N | 02°59'28.71"W | 2008 | 10 |
|  | Cañas, Spain | 42°29'0.09"N | 02°24'17.14"W | 2015 | 10 |
|  | Valpierre, Spain | 42°27'11.00"N | 02°51'48.00"W | 2015 | 10 |
|  | Mateo, Spain | 42°30'34.00"N | 02°44'05.00"W | 2015 | 10 |
|  | Perdiguero, Spain | 42°17'18.78"N | 01°58'50.55"W | 2015 | 9 |
|  | Valbornedo, Spain | 42°24'48.25"N | 02°34'10.06"W | 2015 | 10 |
|  | Villar, Spain | 42°24'03.00"N | 02°42'15.00"W | 2015 | 9 |
| **South-east**<br>**hybrid zone** | Albufeira, Spain | 39°57'0.00"N | 04°14'59.99"E | 2008 | 5 |
|  | Carraixet, Spain | 39°30'06.41"N | 0°19'30.44"W | 2008 | 10 |
|  | Amposta, Spain | 40°38'20.84"N | 0°36'49.32"E | 2008 | 1 |
|  | Marjal del Moro, Spain | 39°37'49.57"N | 0°16'17.14"W | 2008 | 11 |
| **Allopatric**<br>***I. elegans*** | Liverpool, UK | 53°22'57.04"N | 02°56'16.83"W | 2008 | 6 |
|  | Marais D'Orx, France | 43°35'58"N | 01°23'30.00"W | 2014 | 9 |
|  | Saint Cyprien, France | 42°38'23.53"N | 03°01'13.00"E | 2015 | 10 |
|  | Vigueirat, France | 43°37'46.87"N | 04°40'45.24"E | 2014 | 10 |
|  | Leuven, Belgium | 50°51'49.83"N | 04°47'54.65"E | 2015 | 8 |
|  | Kaiserslautern, Germany | 49°23'36.80"N | 07°41'40.89"E | 2008 | 4 |
| **Total** |  |  |  |  | **237** |

**Supplementary Table 2.**
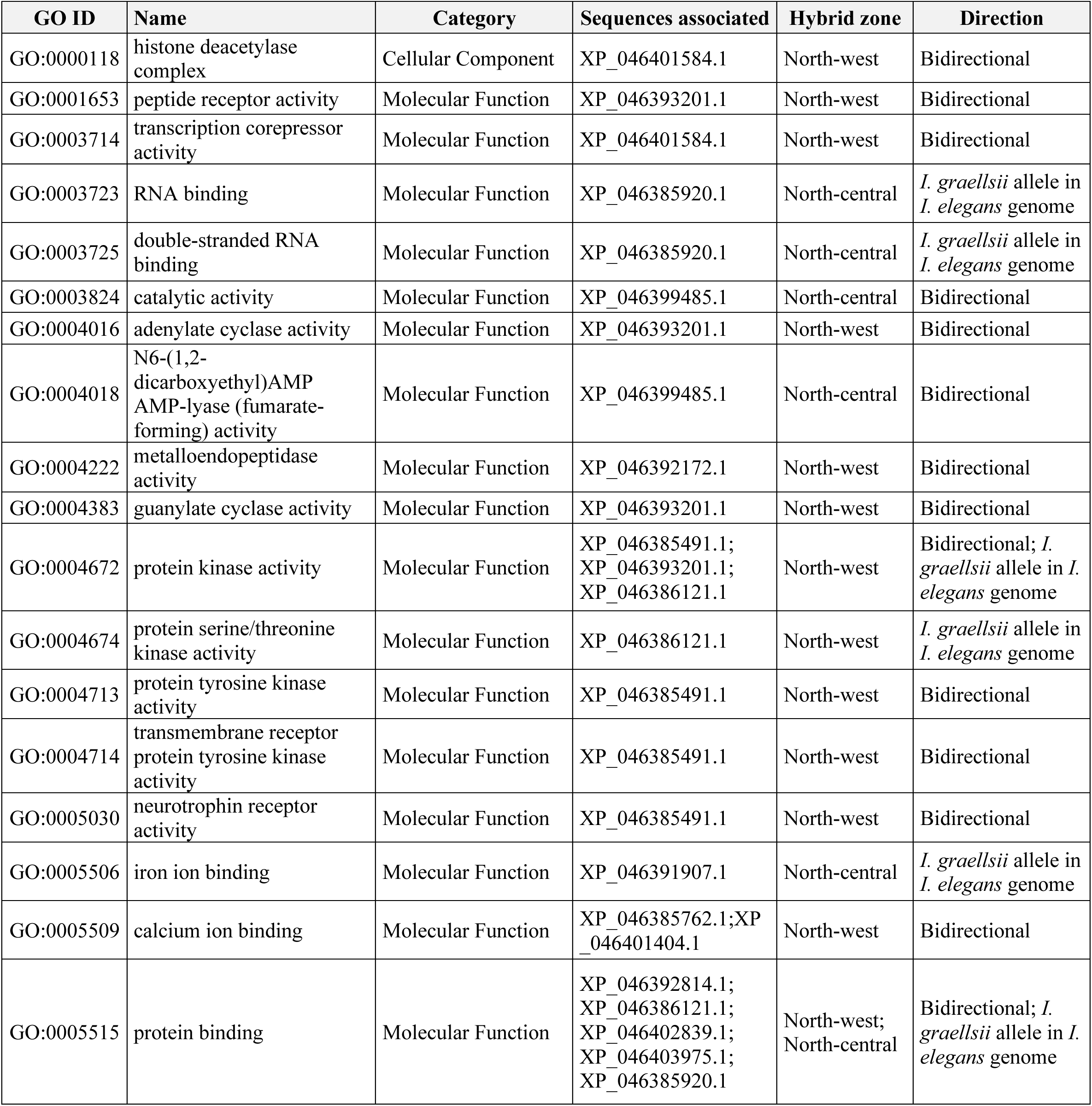
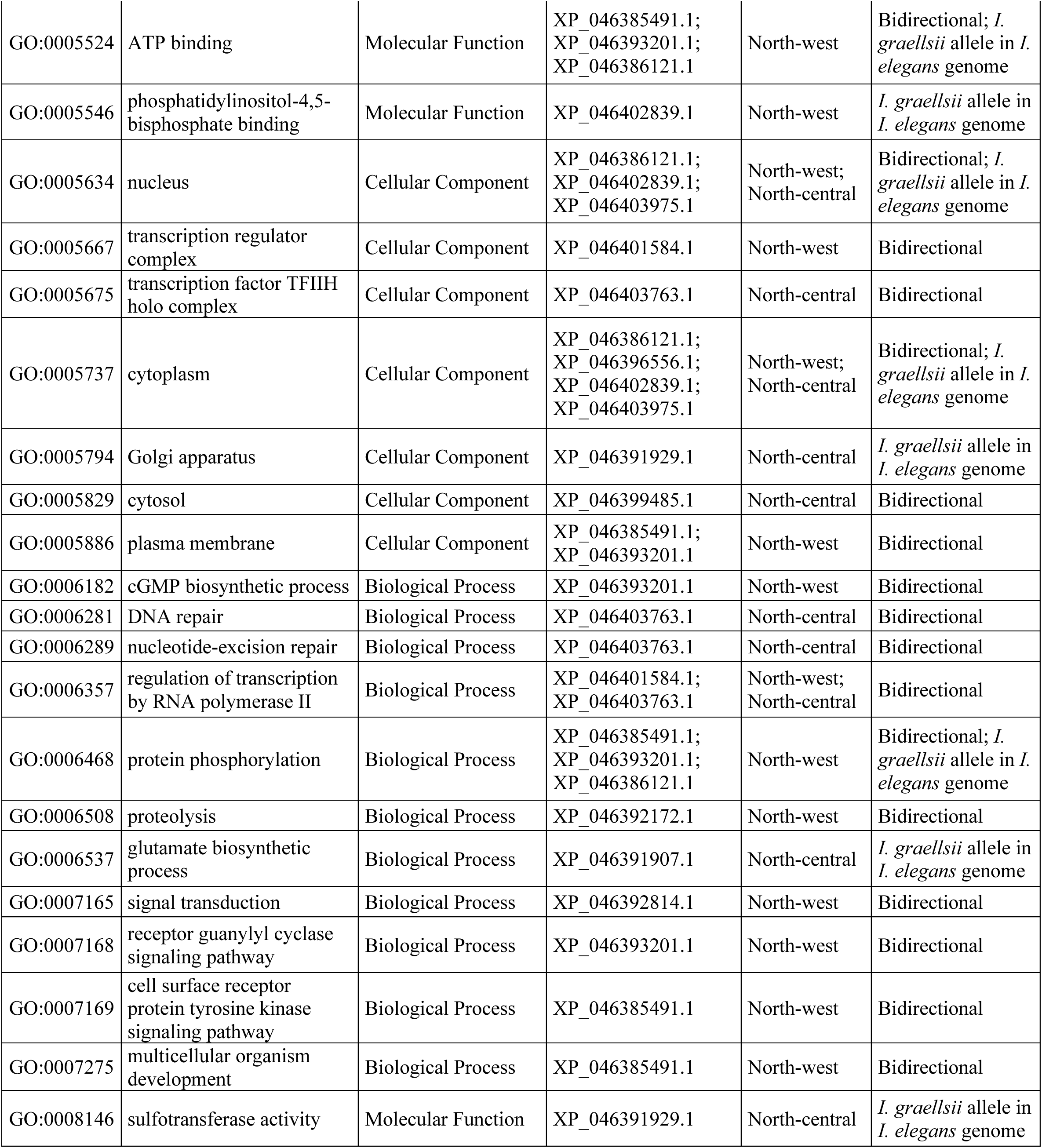
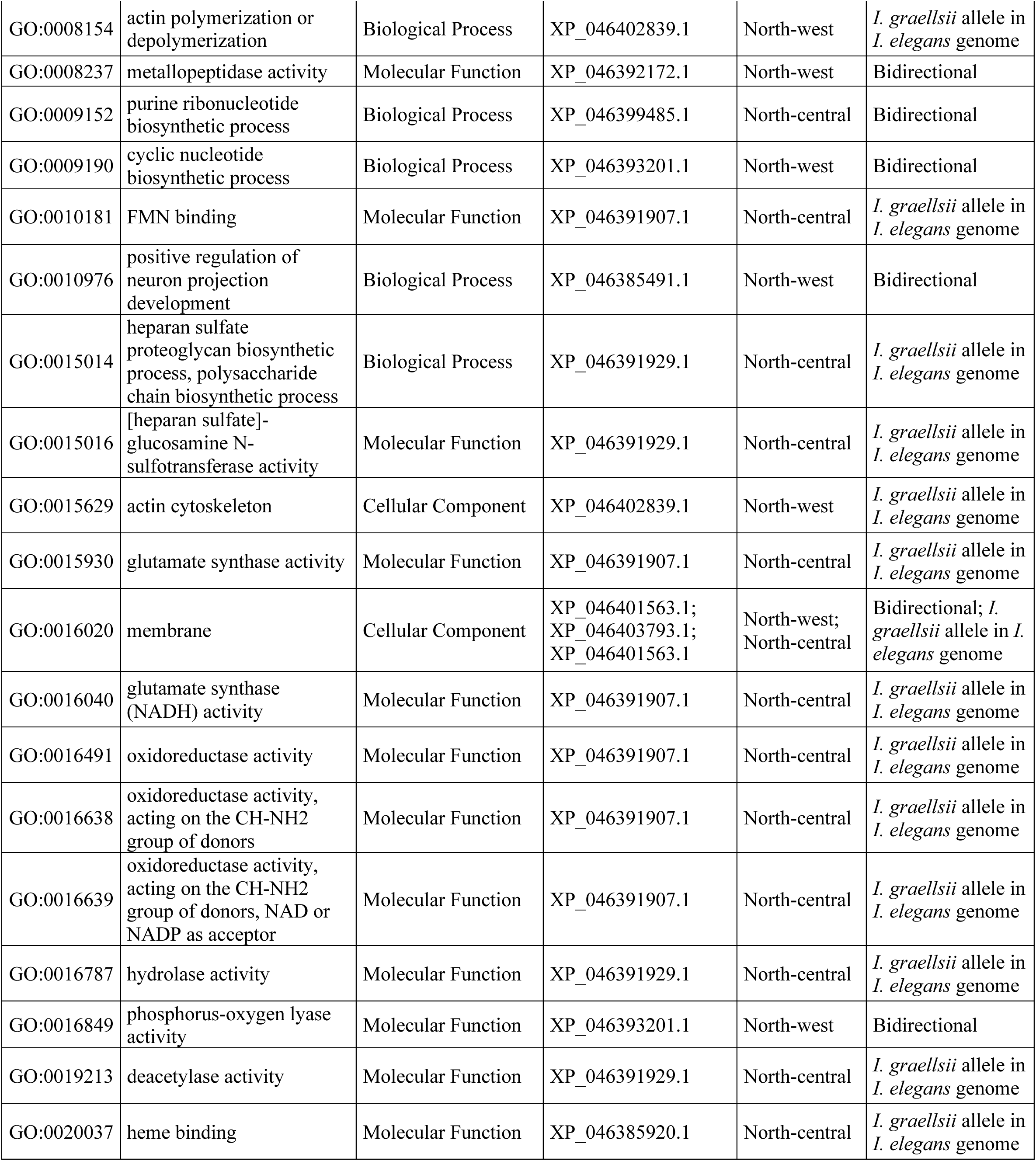
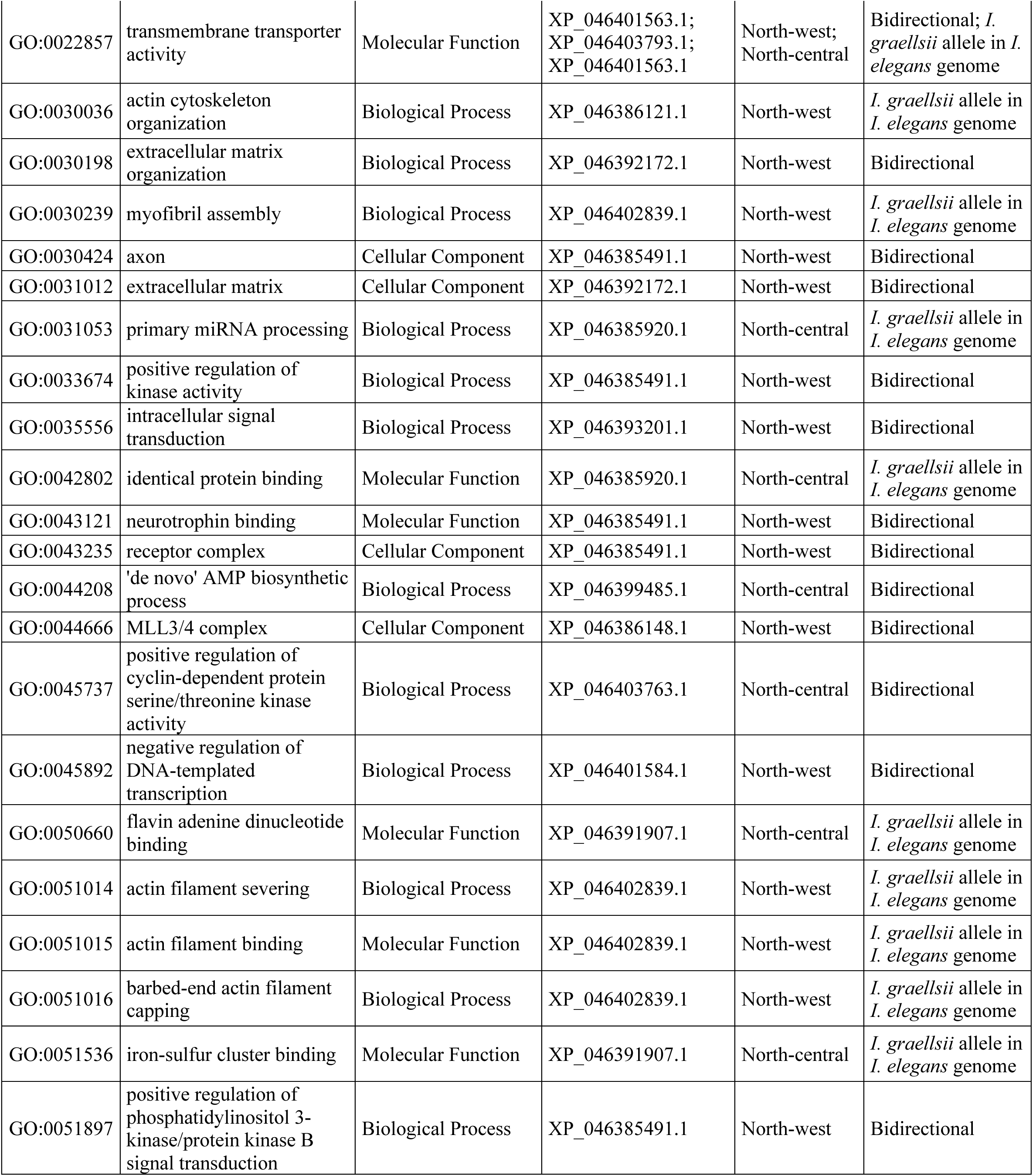
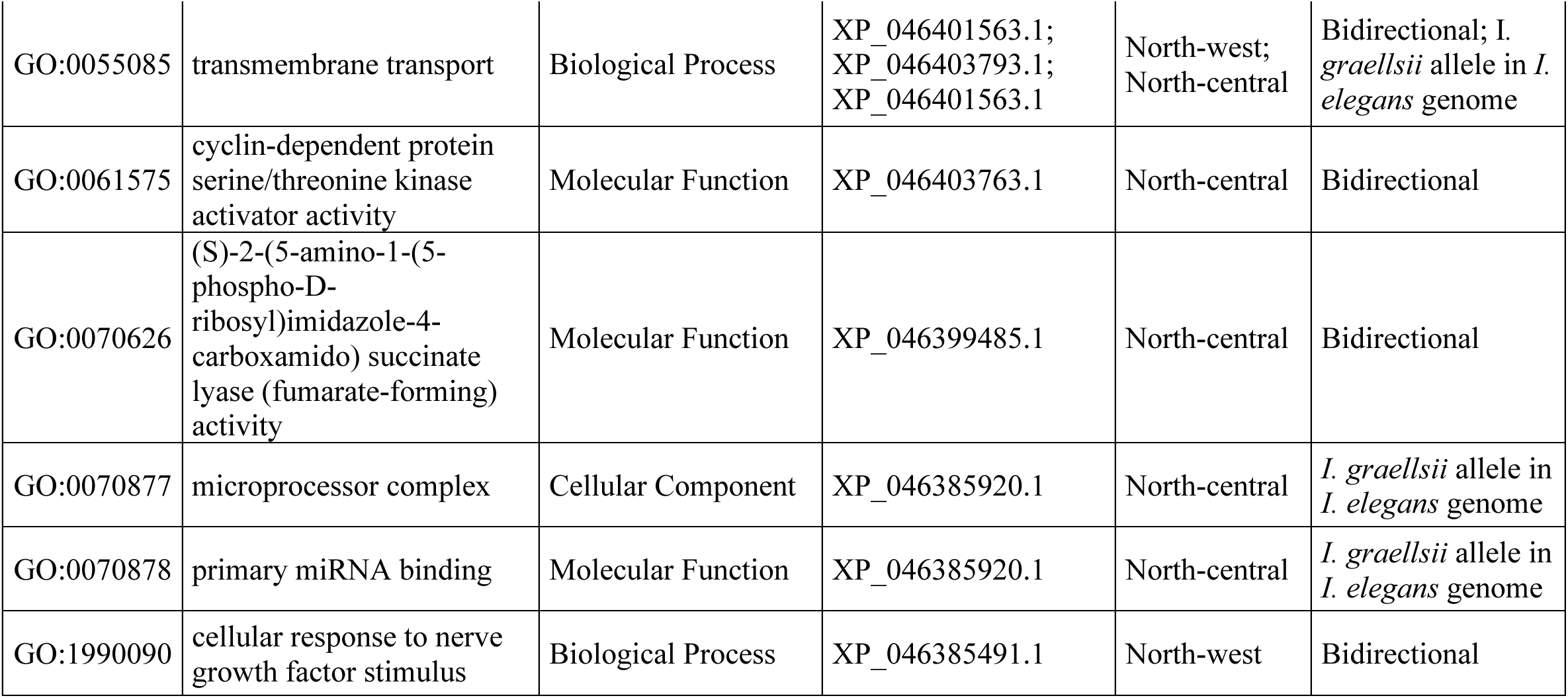
Gene Ontology (GO) terms assigned to protein sequences associated with outlier SNPs. For each term, the table reports the GO identifier, term name, ontology category, associated protein sequences, the hybrid zone where the term was detected, and the direction of introgression.

